# Revealing the genetic components responsible for the unique photosynthetic stem capability of the wild almond *Prunus arabica* (Olivier) Meikle

**DOI:** 10.1101/2021.08.29.458050

**Authors:** Hillel Brukental, Adi Doron-Faigenboim, Irit Bar-Ya’akov, Rotem Harel-Beja, Ziv Attia, Tamar Azoulay-Shemer, Doron Holland

## Abstract

Almond (*Prunus dulcis* (Mill.) D. A. Webb) is a major deciduous fruit tree crop worldwide. During dormancy, under warmer temperatures and inadequate chilling hours, the plant metabolic activity increases and may lead to carbohydrate deficiency. *Prunus arabica* (Olivier) Meikle is a bushy wild almond species known for its green, un-barked stem, which stays green even during the dormancy period. Our study revealed that *P. arabica* green stems assimilate significantly high rates of CO_2_ during the winter as compared to *P. dulcis* cv. Um el Fahem (U.E.F), and may improve carbohydrate status throughout dormancy. To uncover the genetic inheritance and mechanism behind the *P. arabica* **S**tem **P**hotosynthetic **C**apability (SPC), a segregated F1 population was generated by crossing *P. arabica* to U.E.F. Both parent’s whole genome was sequenced, and a single nucleotide polymorphism (SNP) calling identified 4,887 informative SNPs for genotyping. A robust genetic map for U.E.F and *P. arabica* was constructed (971 and 571 markers, respectively). QTL mapping and association study for the SPC phenotype revealed major QTL (log of odd (LOD)=20.8) on chromosome 7, and another minor but significant QTL on chromosome 1 (LOD=3.9). Finally, a list of 73 candidate genes was generated. This work sets the stage for future research to investigate the mechanism regulating the SPC trait, how it affects the tree’s physiology, and its importance for breeding new cultivars better adapted to high winter temperatures.

## Introduction

Almond, *Prunus dulcis* (Mill.) D. A. Webb, is a major fruit tree crop worldwide. As a deciduous fruit tree, it enters dormancy during early winter and renews growth following the fulfillment of a variety-specific period of exposure to low temperatures, known as chilling requirements (CR) and adequate heat requirements. Exposure to a sufficient number of low winter-temperatures is essential for synchronized flowering in the early spring followed by efficient pollination, fruit set and fruit development^1^. CR limit growing areas of deciduous fruit trees and dramatically influence the yield and quality of fruit^2^. When winter temperature increases, CR are not sufficiently provided, and the metabolic activity increases^3^. As a result, carbohydrates are consumed, and intense starch synthesis occur. These changes lead to soluble carbohydrate (SC) deficiency in the buds during the period of flowering and fruit set, which results in disruptive flowering that may reduce yield^4–6^. The ability of the dormant almond to respond this energy depletion is restricted, mainly due to the shortage in photosynthetic leaves during dormancy. Climate changing trends emphasize the urgent need for deciduous fruit crops to gain more plasticity for maintaining their nonstructural carbohydrate (NSC) reserves in warmer winters^2,7^.

*Prunus arabica (Olivier) Meikle*, also known as *Amygdalus arabica Olivier*, defined as a different species from the domesticated almond *P. dulcis*. However, both belong to the *Prunus* genus and are a part of the Rosacea family. The species “ arabica” was named after the geographical region where it was first described. This taxon is native to the temperate-Asia zone. It covers the Fertile Crescent Mountains, Turkey, Iran and Iraq. In the Middle East it can be found in Lebanon, Syria, Israel (Judean Desert) and Jordan^8^. *P. arabica* can be found in altitudes between 150-1,200 m and rarely up to 2,700 m. It is a bush, rather than a tree, with a very long root system and is considered resistant to drought^9,10^. As a deciduous tree, *P. arabica* drops its leaves at the end of the summer, turns meristems into buds and stops growing. However, unlike other almond species, it’s young branches remain green and are not covered with bark (i.e., no cork layer deposition) throughout the dormancy phase (Fig. 1 a-d). In fact, *P. arabica* stems remain moist and green during the whole year. *P. arabica* green stems were previously suggested to photosynthesize^11^, yet no physiological evidence was published regarding their ability to assimilate external CO_2_.

**Figure 1.**
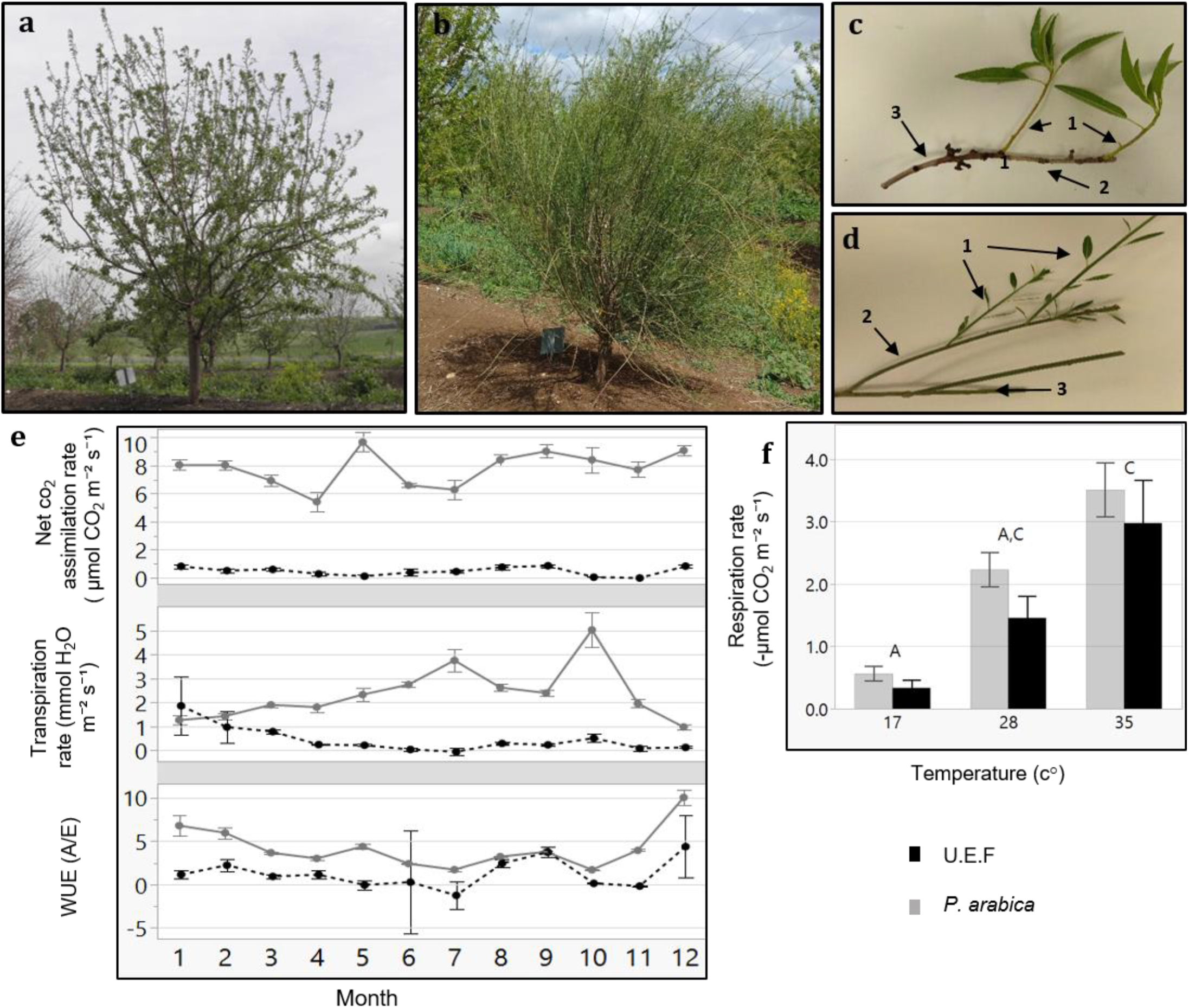
Gas exchange measurements of *P. arabica* and *Prunus dulcis* cv. Um el Fachem (U.E.F.) stems. Three years old trees of the Israeli cultivar *Prunus dulcis* cv. U.E.F (a), and the wild almond *P. arabica* (b) at spring. Stems of U.E.F. (c), and *P. arabica* (d) annually developed: one-year old (c1, d1), second year (c2, d2) and three years old stems (c3, d3). Gas exchange data of one-year old stems along the year (e) of *P. arabica* (solid gray line), and U.E.F (dashed black line). Each dot denotes the average of two independent days of measuring for each month (n=8). Stem respiration rate in response to three different temperatures (f) of *P. arabica* (grey bar) and U.E.F (black bar). Four stems were measured for each genotype in each temperature (n=4). Different capital letter represents significance (α=0.05) between temperatures, not between species. The error bars represent ± SE.

Stem photosynthesis was previously shown in other desert species^12^. In these species, which do not belong to the Rosacea family, high efficiency of CO_2_ assimilation comparable with that of the leaf was demonstrated. Because stem photosynthesis was found in desert plants, it was suggested that it might play a role in carbon gain under stress conditions such as heat and drought^13^. The contribution of stem photosynthesis to tree adaptation under drought was further supported by evidence showing that stem photosynthesis assists to embolism repair^14,15^. Ability to photosynthesize through stems could prove to be highly beneficial for deciduous fruit trees such as almonds in maintaining the energy balance of the tree, particularly during hot winters and springs when respiration is enhanced, and tree energy is limited due to leaf drop and the lack of photosynthetic organs ^7,16^. Previous genetic studies of *P. arabica* were limited to phylogenetic studies and encompass a small number of markers (few to dozens)^17,18^. To the best of our knowledge, no genetic approach detected to uncover the mechanism of the stem photosynthesis phenomena.

Recent important advancement in Rosacea genetics and genomics enables the application of a genetic approach in the study of important physiological processes. Such advancements include the development of genetic maps based on F1 and F2 populations^1^, and their usage for mapping QTLs affecting CR in apple (*Malus domestica*)^19^, pear (*Pyrus communis*)^20^, apricot (*Prunus armeniaca)*^21^, and peach (*Prunus persica)*^22^. In almonds, genetic mapping of hybrid populations based on distinctive CR, demonstrated that a major gene, *LATE BLOOMING* (*LB*) was associated with blooming date and dormancy release^23^. In addition, complete genomes and various transcriptomic datasets of Rosacea: apple, cherry *(Prunus avium)* and peach^24^ were published. Recently, two almond genomes were published: *P. dulcis cv*. Texas (https://www.ncbi.nlm.nih.gov/bioproject/572860) and *P. dulcis cv*. Lauranne (https://www.ncbi.nlm.nih.gov/bioproject/553424). This trend of new genomic and transcriptomic data of deciduous fruit trees, including almonds, sets the stage for intense genetic research and the development of novel marker-assisted breeding approaches. In this report, we combined physiological and genetic approaches to study the green stems of *P. arabica* throughout the year. By direct measurements of gas exchange, we demonstrate that *P. arabica* stems transpire and assimilate significant levels of CO_2_ all year round, including during the dormancy period. Moreover, we undertook a forward genetic approach, and used an F1 population segregating for the stem photosynthesis trait for high-resolution mapping of major QTLs for this trait. The genetic markers and candidate genes that our study underlines pave the way to undermine the physiological role of stem photosynthesis and its utilization for genetic improvement.

## Results

### P. arabica assimilate CO_2_ through green stems

*P. arabica* stems remain green during winter while cultivated almonds develop an outer grey cork layer (Fig.1b, d). To study if these stems are actively assimilating CO_2_, we undertook gas exchange measurements of tree stems in the orchard with the Licor 6800 Portable Photosynthesis System. Two different almond species were compared, the wild almond *P. arabica* and the cultivated almond *P. dulcis* (U.E.F), throughout the entire year (Fig. 1e). The data indicate that *P. arabica* assimilates CO_2_ through its green stems during all year (annual average of 8±0.19 µmol CO_2_ m^-2^ sec^-1^), while similar one-year old stems of U.E.F assimilation capacity is almost nil (annual average of 0.5±0.05 µmol CO_2_ m^-2^ sec^- 1^). The significantly high CO_2_ assimilation rates of *P. arabica* stems were found comparable with assimilation rates of *P. arabica* leaves (11.2±0.8 CO_2_ m^-2^ sec^-1^, July average, data not shown). Although some fluctuations were observed between the different seasons, pronounced high CO_2_ assimilation rates were found in *P. arabica* stem during the whole year (Fig. 1e). Finally, *P. arabica* stem transpiration rate is relatively low in the dormancy phase and gradually increases until it peaks in October (1.2±0.18 in January to 5±0.7 mmol H_2_O m^−2^ sec^−1^ in October). In contrast, transpiration from U.E.F stems is relatively constant and low throughout the year (0.46±0.11 mmol H_2_O m^−2^ sec^−1^; Fig. 1e). Transpiration rate fluctuation also attribute to high instantaneous water use efficiency (iWUE) of *P. arabica* during the dormancy phase (two-fold higher than U.E.F in December; Fig. 1e).

Previous studies on temperate fruit trees showed a positive correlation between tissue temperature and SC consumption through respiration^16^. To find out how stem respiration of *P. arabica* and U.E.F are influenced by temperature, we measured the respiration rate of one-year old stems while exposing them to three different temperatures (17°, 28° and 34° C) (Fig. 1f). Increased respiration rate in response to elevated temperature was observed in both almond species (0.5±0.11, 2.2±0.27, 3.5±0.44 and 0.33±0.12, 1.4±0.35, 3±0.68 µmol CO_2_ m^-2^ sec^-1^ for *P. arabica* and U.E.F respectively for each temperature), while no significant differences were observed between species (for each measured temperature).

### Stem assimilation is genetically inherited

To elucidate the genetic nature of the assimilating stem trait of *P. arabica*, an F1 hybrid population (n = 92) was established by crossing *P. arabica* (male) with U.E.F (female) (Fig. 2c, d). The same approach of gas exchange measurements in the field was used for phenotyping the SPC trait among the three year-old F1 population during dormancy. Twelve offspring assimilated CO_2_ via their stems in a similar level as *P. arabica* (offspring 24H27 is the highest; 8.3±0.14 µmol CO_2_ m^-2^ sec^-1^), and Thirty-seven individuals assimilated as U.E.F or less (Fig. 2a). Analysis of distribution demonstrated two prominent peaks within the histogram (Fig. 2b). Although the ‘3 Normal Mixture’ is the most accurate to describe the phenotype distribution (achieved the lowest AICc and the -2Log Likelihood values), broad-sense heritability (h^2^) was found to be high (0.91).

**Figure 2.**
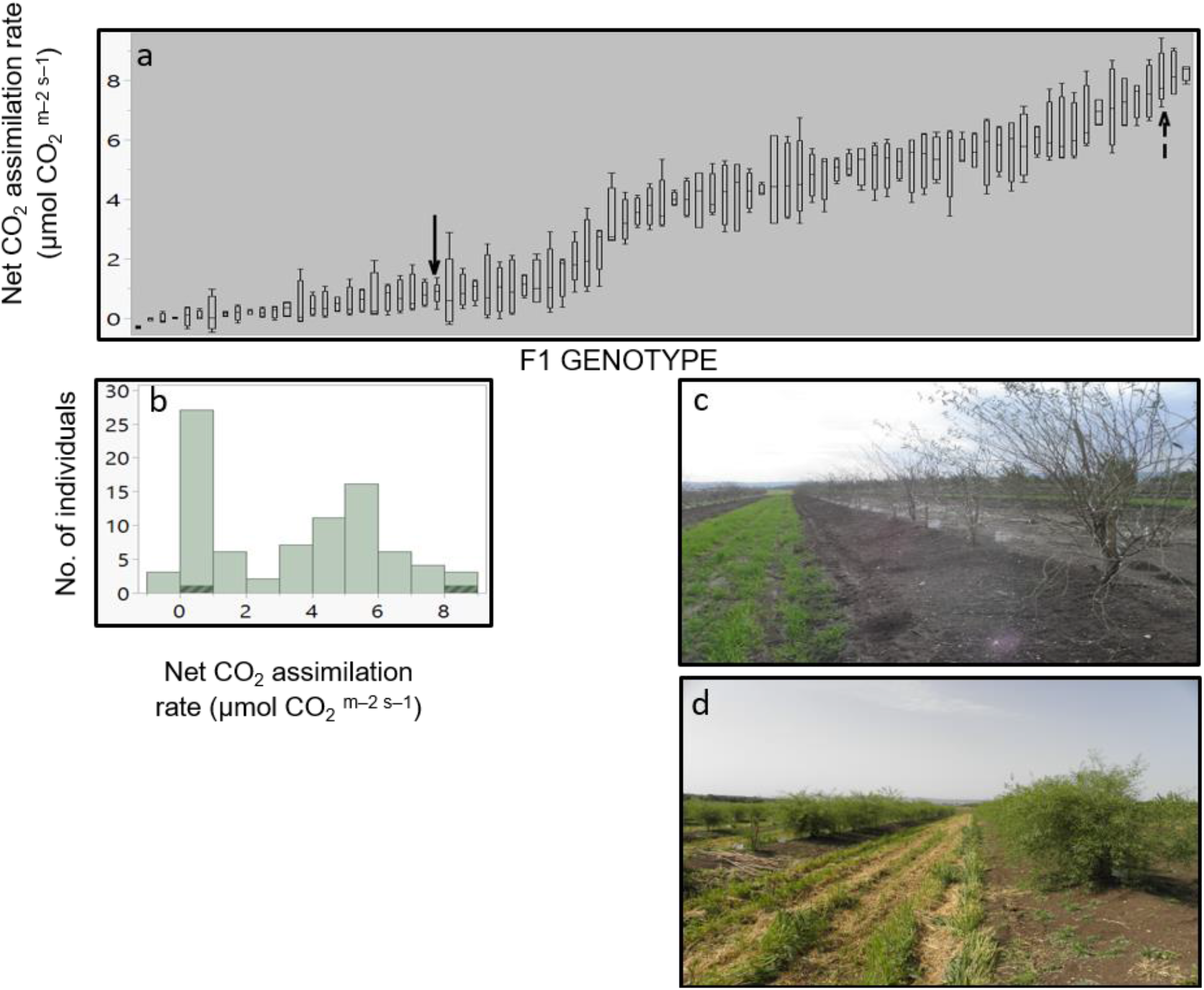
Stem photosynthetic capability (SPC) in the F1 progeny. Levels of net CO_2_ assimilation for each offspring (a). Each box plot presents the average of four stems (n=4). The F1 progeny parents, *P. arabica* and U.E.F are marked by dashed arrow and simple arrow, respectively. Distribution histogram of the same data is presented in (b), while parents’ data is highlighted. Measurements were conducted during February 2020 while the trees were dormant. Representative pictures of the F1 population while dormant (c) in February, and during the vegetative phase in April (d).

### Sequence comparisons, SNPs identification, and genotyping of the F1 population

Segregation of the SPC trait among the F1 population rendered it as suitable for genetic mapping. For this purpose, we sequenced the *P. arabica* and the U.E.F genomic DNA, targeting for high coverage, to ensure reliable (SNP) calling. The reads were aligned against the reference genome of *P. dulcis cv*. Lauranne, which was found as the closest (> 97% mapped reads) of the two published almond genomes (Table 1).

**Table 1.** Quality data from whole genome sequencing of *P. arabica* and U.E.F. Quality parameters (Q20, Q30) from sequencing of each parent is presented with respect to the reference genomes of *P. dulcis* cv. Lauranne and *P. dulcis* cv. Texas. The row sequence data was mapped to each of the reference genomes. The total variant sites (SNPs and InDels) between each species and cv. Lauranne reference genome are also presented.

| Species | Average coverage | Q20 | Q30 | % Mapping VS Lauranne | % Mapping VS Texas | Total variant sites | % Heterozygous variant sites |
| --- | --- | --- | --- | --- | --- | --- | --- |
| *P. arabica* | ~ X 57 | 94.5 | 87.37 | 97.93 | 85.94 | 3,750,363 | 35.18 |
| U.E.F (*P.dulcis*) | ~ X 55.5 | 94.7 | 87.71 | 98.37 | 89.82 | 2,407,787 | 69.29 |

A total of 3,750,363 and 2,407,787 variants (i.e., SNPs or short InDels) were detected for *P. arabica* and U.E.F respectively, against the cv. Lauranne reference genome. Analyzing the variants showed that 71.5% and 72.6% (*P. arabica* and U.E.F, respectively) are in the intergenic region (Table S1). Furthermore, a higher variant number was detected in the intronic regions in relation to the exons (Table S1). The initial number of identified SNPs (Total variant sites in Table 1) were filtered by several types of criteria as specified in materials and methods.

Overall, 4,887 SNPs that are heterozygous for one of the parents and homozygous for the second were selected for F1 genotyping screening. The SNPs are spread at intervals of about 40K along the almond genome.

The F1 population was successfully genotyped with 4,6125 SNPs. The resulting genotyping quality data (Table S2) represent high coverage (152X), and low number of missing data (5.5%). Further analysis of the genotyped F1 population with the SNP panel described above show the allelic frequency within the F1 population is 50%, as expected from an F1 population (Fig. 3). However, since the allelic composition in this bi-parental population is AA x Aa, we can also refer this ratio as the allelic frequency of the heterozygous genotype. Therefore, data presented (Fig. 3) also indicates exceptional chromosomal regions (hot spot) with a unique pattern of inheritance that deviates from the 1:1 ratio, for example, in chromosome 3 (see black arrow in Fig. 3).

**Figure 3.**
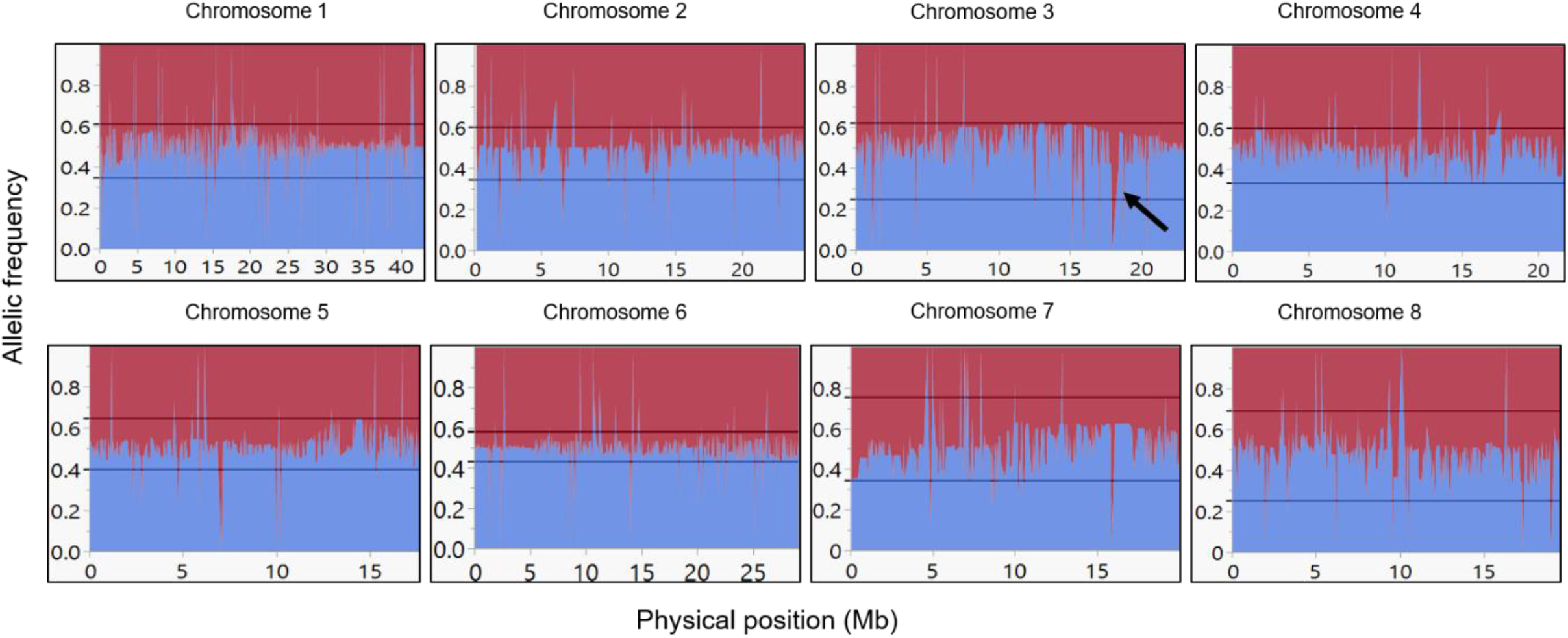
Allelic frequency among the F1 population. Allelic frequency of the heterozygous allele in the F1 population for each marker. Blue line indicates the *P. arabica* allelic incidence, and red line indicates the U.E.F allelic incidence. X axis is the physical position in Mb, and Y axis represents the allelic ratio (from 0 to 1). Black arrow in chromosome 3 represents an example of un-expected deviated region (“ hot spot”). References lines presented the ‘tolerance interval’ limits.

### Construction of genetic maps for the F1 population

To establish a genetic map of the F1 population, Join Map 4.1 software was used^25^. CP (cross pollination) population type was performed with the lmxll code for markers that were homozygous for the male parents (*P. arabica*) and heterozygous for the female parent (U.E.F). The code nnxnp used for the opposite case. A significant portion of the markers was filtered, most of them due to complete similarity (∼50%). Overall, 1,533 SNPs were used for mapping (Table 2). Because there were no common markers for both parents, the hkxhk code was not applied. Using the pseudo test cross method^21^, we separated the markers for two different maps: one map for the U.E.F (where *P. arabica* is homozygous, lmxll code), and the second map for the *P. arabica* parent (where U.E.F is homozygous, nnxnp code). Applying this strategy, we obtained two maps with robust numbers of markers and good density. The U.E.F map was found to be denser than the *P. arabica* map and includes 971 markers with an average distance of 0.533 centiMorgen (cM), while *P. arabica* map contains 572 SNPs with an average distance of 1.093 (cM) (Table 2). It can be clearly seen that the distribution of the SNP markers is well spread over the eight almond linkage groups (LG) (Fig. 4).

**Table 2.**
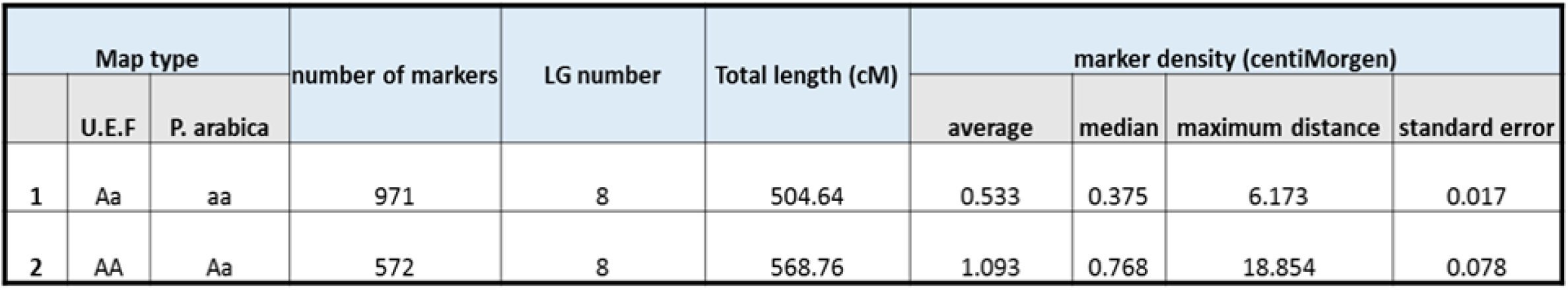
Characteristics of the established genetic maps. U.E.F map (1) was generated from markers that were homozygous in *P. arabica* (male) and heterozygous in the U.E.F (female). *P. arabica* map (2) was generated from markers that were homozygous for U.E.F and heterozygous for *P. arabica*.

| Map type |  |  | number of markers | LG number | Total length (cM) | marker density (centiMorgen) |  |  |  |
| --- | --- | --- | --- | --- | --- | --- | --- | --- | --- |
|  | U.E.F | *P. arabica* |  |  |  | average | median | maximum distance | standard error |
| 1 | Aa | aa | 971 | 8 | 504.64 | 0.533 | 0.375 | 6.173 | 0.017 |
| 2 | AA | Aa | 572 | 8 | 568.76 | 1.093 | 0.768 | 18.854 | 0.078 |

**Figure 4.**
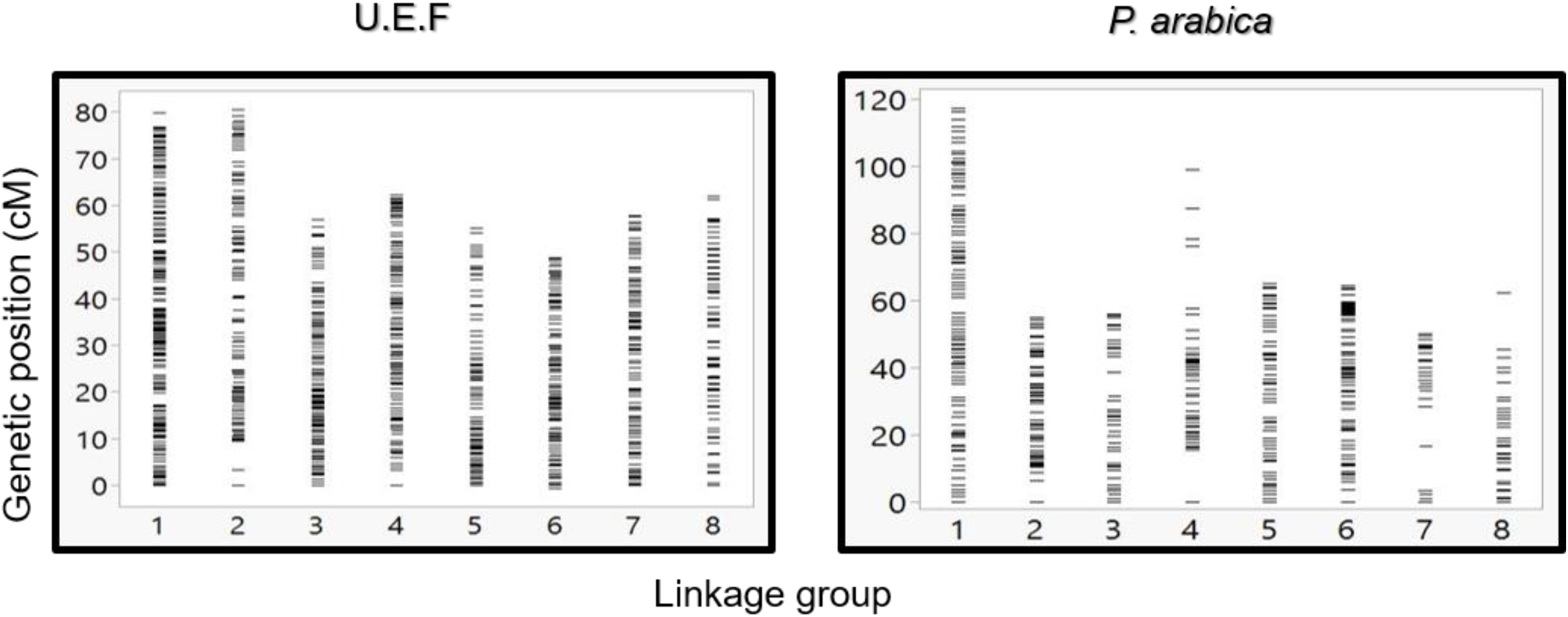
Graphic presentation of markers density and distribution along the eight linkage groups. Comparison between U.E.F map (left graph) and *P. arabica* map (right graph). Each horizontal line represents a single marker.

To assess the validity of the genetic map, the order of SNP markers as determined by the U.E.F genetic map was compared with the deduced order from the physical map as determined by cv. Lauranne reference genome. The analysis (Fig. 5) demonstrates a good co-linearity between the genetic and the physical map. Remarkably, most of the markers from the genetic map were highly correlated with the physical order (Fig. 5), yet, few markers did not correlate (chromosome 6, Fig. 5). The genetic map divided the markers into eight LGs parallel to the previously published chromosome organization order. Moreover, the slopes generated between the physical orders to the genetic order represent recombination frequency (cM / Mb). Thus, one can see that around the centromere, the slope is more horizontally, meaning the cM / Mb ratio is relatively low. Thirty-eight SNP markers representing un-scaffold contigs (i.e., chromosome 0) in the reference genome project were assembled into six linkage groups based on the genetic maps (marked by yellow dots in Fig. 5).

**Figure 5.**
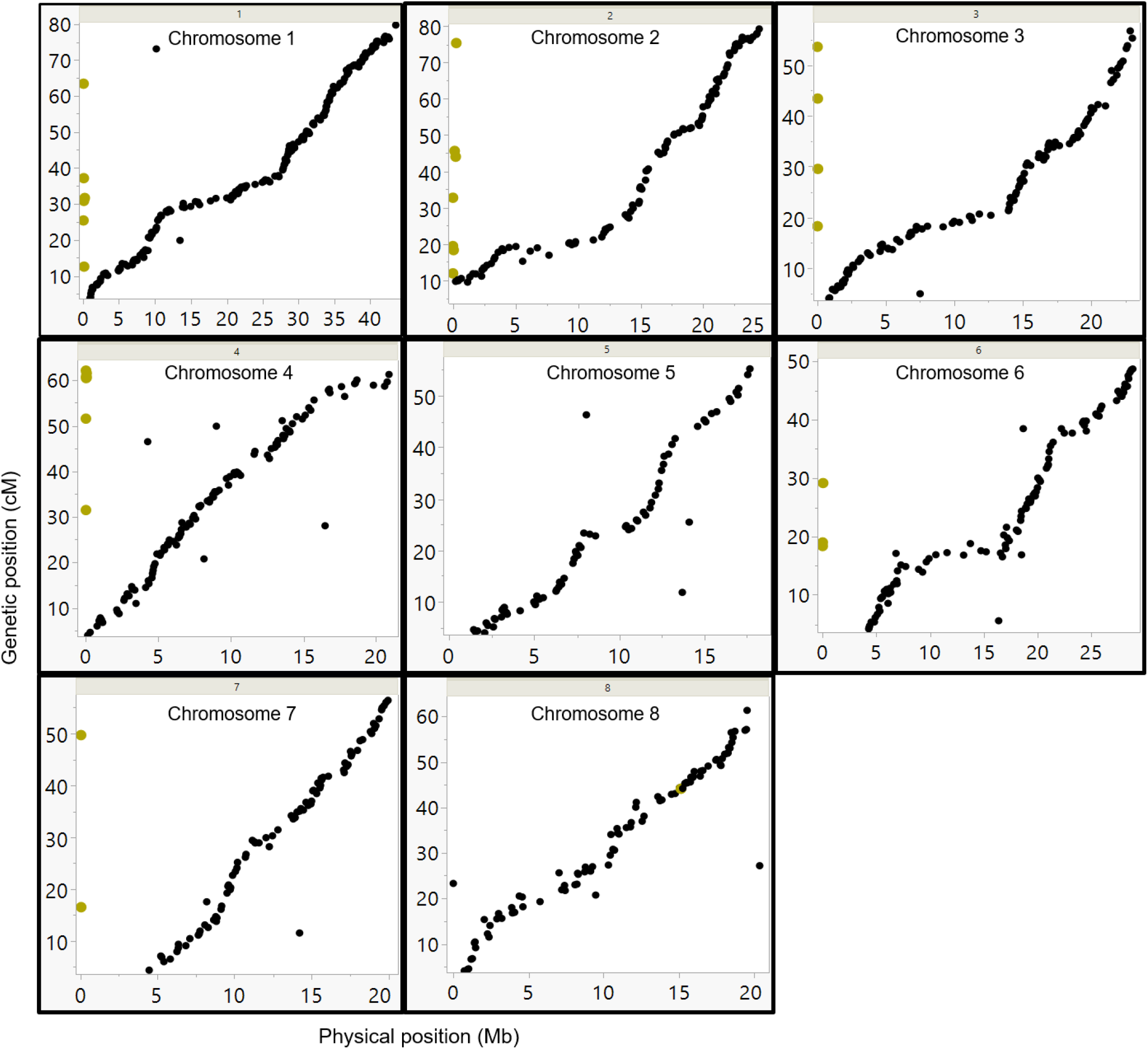
Comparison between the genetic and physical order of the markers. SNP markers were placed according to their physical position on the Lauranne reference genome sequence (X axis), and their position on the U.E.F genetic map (Y axes). The yellow dots represent markers from unplaced scaffolds, according to the Lauranne reference genome (chr-0). Those markers were mapped to several chromosomes in this study based on the genetic map data.

### QTL analysis and genome wide association study (GWAS) of the SPC trait

Two main approaches were initiated for detecting genomic regions regulating the SPC. QTL mapping, computed with Map QTL by interval mapping (IM) analysis^26^ and Genome wide association (GWAS) by TASSEL software^27^. QTL mapping generated two significant QTLs. Each QTL was discovered only in one of the two genetic maps. Thus, one major QTL (LOD=20.8) was mapped on LG 7 spanning a region of 2.4 cM detected on the U.E.F map. The second, minor but significant QTL (LOD=3.9) was detected at the end LG 1, spanning a region of 4.4 cM on the *P. arabica* map (Table S3, Fig. 6).

**Figure 6.**
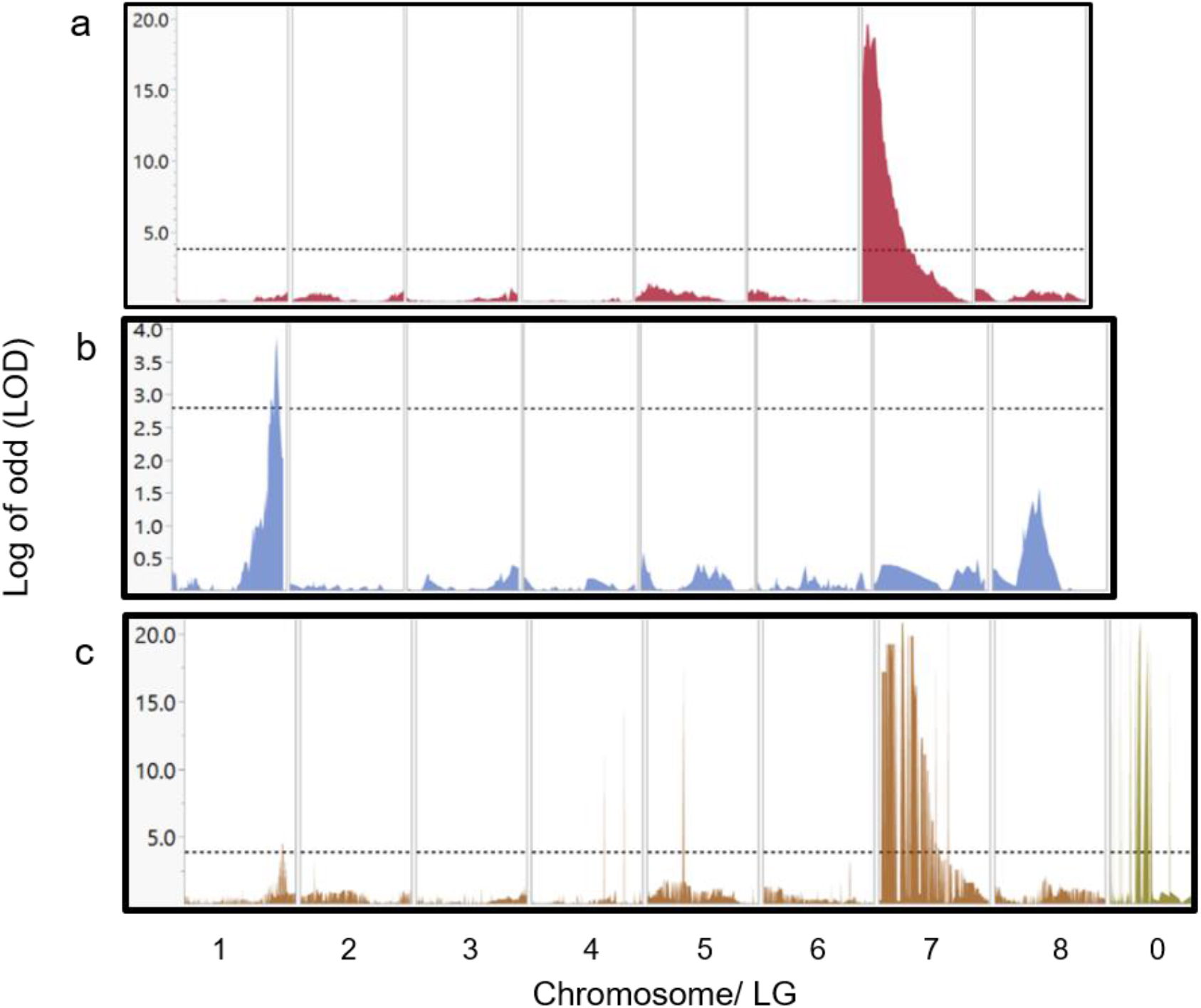
QTLs and GWAS analysis for the photosynthetic stem trait. Major QTL of 2.4 Cm width and LOD score of 20.8, was detected in chromosome 7 by using the U.E.F genetic map (a). Minor QTL of 4.4 Cm width, and LOD score (3.9) was located at the end of chromosome 1 by using the *P. arabica* genetic map (b). Results of GWAS using the whole set of markers (3,800) sorted by their physical position according to the reference genome (c), revealed both loci in chromosome 7 and chromosome 1. Markers in c were sorted by their physical position according to the reference genome. Markers that were placed on chromosome 0 and found as highly associated to the SPC trait are also shown in (c). The horizontal dash line represents significance level according to permutation test (1000 times at α=0.05).

Applying GWAS approach with TASSEL enabled us to simultaneously detect two genomic sites that regulate the SPC on chromosomes 1 and 7 at positions similar to those detected by QTL mapping. The major region on chromosome 7 spanning only 400kb, and the minor on chromosome 1 containing 700kb. Moreover, GWAS analysis showed significant associations with markers aligned to chromosome 0. Interestingly, two of these markers assembled into the major QTL in locus 7 (Table S3, marked with gray background). The major QTL explained 67% of the phenotypic variance, while the minor QTL explained 19.3% (Table S3).

### QTL’s interaction

As presented, two significant loci were discovered as regulating the SPC (Fig. 6, Table S3). Full factorial test shows a significant additive effect between those two associated loci (<0.0001; Fig. 7a, b). Yet, no epistatic effect was found (p-value= 0.676; Fig. 7c). As expected, in both QTLs the *P. arabica* alleles were the increasing alleles regarding the SPC trait.

**Figure 7.**
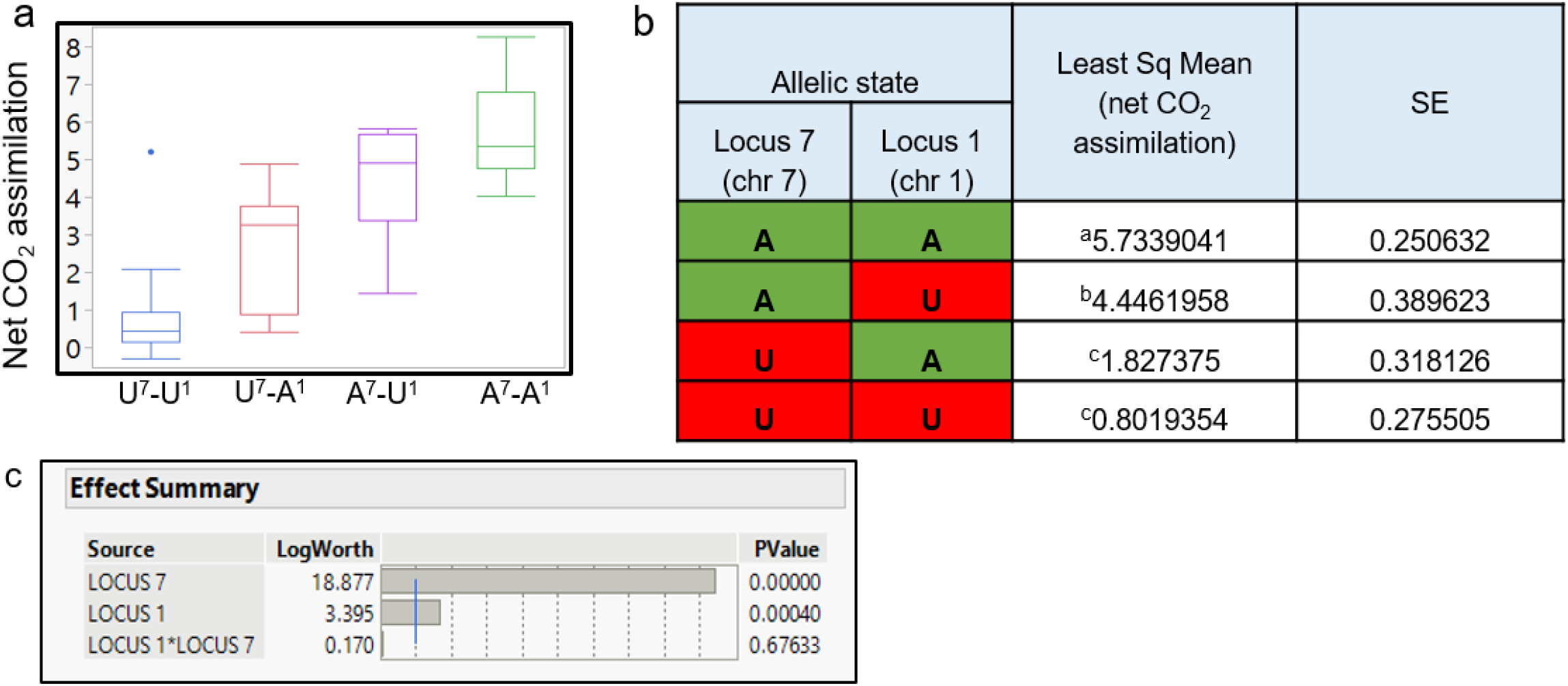
QTLs effect on net CO_2_ assimilation and the synergistic effect between QTLs. Least square of means for each allelic combination is presented (a). Each box plot represents the population individual’s average phenotype-grouped for their allelic combination. Y axis is the level of net CO_2_ assimilation. The X axis represents the allelic combination. On the X axis, the capital letter A, refers to individuals with *P. arabica* allele combination, U refers to individuals with U.E.F allele combination in each one of the QTL, and superscript numbers (7, 1) present the QTL identity. Numerical presentation of the data is presented in a (b). *P. arabica* allele combination is marked in green, and U.E.F marker combination is marked in red. Different letters indicate significance (α=0.05). Statistical evaluation of each QTL effect, and interaction between the two loci (c). Presented results were analyzed by the “ Full factorial test”, blue line is equivalent for p-value of α=0.01.

### Generating a list of candidate genes

Combining GWAS and QTLs data described above with that of the cv. Lauranne reference sequence allowed us to delineate a list of genes within the regions that are predicted as responsible for the SPC trait. The region at chromosome 1 includes 113 annotated genes with SNPs between *P. arabica* and U.E.F (Table 3). Among those, 17 include non-synonymous SNPs in the genes coding region. The associated region at chromosome 7 consists 336 genes with SNPs, of which only 54 have non-synonymous SNPs in their coding region (Table 3).

**Table 3.**
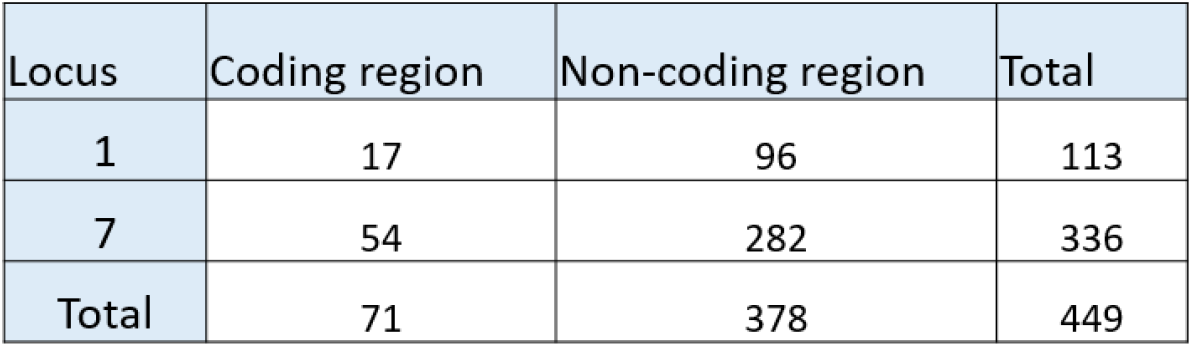
Summary of candidate genes’ list. The annotated genes within the regions spanning the two QTLs are presented. Only genes that have non-synonymous variants between the two parents are presented. Genes are divided for polymorphism region as determined by the D.N.A sequence.

| Locus | Coding region | Non-coding region | Total |
| --- | --- | --- | --- |
| 1 | 17 | 96 | 113 |
| 7 | 54 | 282 | 336 |
| Total | 71 | 378 | 449 |

## Discussion

### P. arabica can assimilate CO_2_ through its green stems all year round

This study investigated the unique wild almond *P. arabica*, a member of the Rosacea family. We demonstrated for the first time that the wild almond *P. arabica* is assimilating external CO_2_ via its green stems all year round, including during the dormancy period when the tree sheds its foliage. In contrast, the cultivated almond U.E.F is unable to do so (Fig. 1e). Interestingly, even the young green stems of U.E.F (Fig. 1c), which are not covered with bark layer, in spring, are assimilating CO_2_ in a negligible levels (Fig. 1e). *P. arabica* is a desert plant, and its native growing ecosystems encompass the desert margins of the Fertile Crescent^8^. Previous studies have indicated that perennial desert shrubs, such as *Ambrosia salsola* and *Bebbia juncea*, are able to photosynthesize through their stems^28^. This capability was attributed to better adaptation of the plant to arid climate conditions^12^. The geographical distribution of *P. arabica* suggests that stem photosynthesis might be important for its survival in dry areas^13^. Indeed, several studies have suggested that stem CO_2_ assimilation may help prevent embolism damage in response to drought^14,15^. Others have claimed that SPC enables more efficient carbon gain with respect to water loss (transpiration) due to smaller surface area when compared to leaves ^29^. Interestingly, fluctuations were observed in transpiration rates of *P. arabica* stem during the year, which resulted in maximal iWUE in winter time since CO_2_ assimilation was not highly affected (Fig.1e). Further physiological and anatomical analyses are needed to understand better this unique SPC trait and its contribution to almonds in harsh climates.

Among the wild and cultivated species of almond, only *P. arabica* and its very close relative, *Prunus scoparia* (Spach) Schneider, are known to produce all-year green stems^30^. Yet, stem CO_2_ assimilation in *P. scoparia* was not demonstrated. To our knowledge, *P arabica* is the only known deciduous fruit tree that possesses the SPC trait. As such, *P. arabica* is unique among all deciduous fruit trees in its capability to assimilate external CO_2_ from stems in winter. In addition to the reasonable assumption that SPC contributes to plant survival in dry conditions by preventing embolism damage and by improving carbon gain, it is anticipated that SPC in winter could influence other critical physiological processes that are dependent on carbohydrate management, including dormancy break and ability to support heavy yield^6,31^. This is particularly true for deciduous fruit trees under the pressure of energy shortage when the trees have to support both leaf and flower development^5^. Deciduous fruit trees rely on their energy reserves during this growth phase since their leaves are not yet fully developed^16^. Recent studies on deciduous fruit trees demonstrate how higher temperatures are positively correlated with tree NSC consumption. It was shown that during higher winter temperatures, NSC reserves depleted rapidly^7^. For setting these results in *P. arabica* and U.E.F, we measured respiration rate in response to elevated temperatures. As expected, environmental temperature indeed raised the respiration rate in both species (Fig. 1f). This result emphasizes the possible link between winter temperatures and trees’ NSC status during winter. Importantly, it highlights the advantage of a functional photosynthetic organ in warm winters during dormancy phase. Nevertheless, to establish the link between tissue temperature to tree NSC and their contribution to yield, we are currently conducting additional research.

### The SPC trait depict a high heritability value and segregates in an F1 population

As a means of investigating the functional role of SPC in almonds, we undertook a forward genetics approach. Identification of the genetic components underlying SPC is crucial for deciphering the role of SPC in respect to dormancy break and adaptation to upcoming climate changes. Moreover, molecular markers for the genes in question could provide an efficient tool for breeding and genetic manipulation for better almond trees. For this purpose, we established an F1 population from a cross between *P. arabica* and U.E.F. Remarkably, SPC measurements of individual progenies within the F1 population indicate that the SPC trait is inherited and segregated already in the F1 population (Fig. 2a). Interestingly, the distribution of the trait is not normal but displays two main peaks, suggesting that this trait is probably controlled by a small number of genes or QTLs (Fig. 2b). Altogether, the data indicated that the F1 hybrid population could be used as a mapping population for the SPC trait. Furthermore, the high heritability value (0.91) suggesting the SPC trait can be integrated into cultivars through classical genetic crosses. Yet, current data is not sufficient to determine the heritability control of the trait (dominant/recessive) due to the continuous nature of the SPC phenotype.

### Genotyping

In this study, we report the feasibility of using a novel methodology to obtain genetic markers through QTL mapping of almond. The method of ‘targeted SNP seq’ that was already described for other plants^32^, takes advantage of the availability of almond chromosomal organization and genome sequencing data. Only markers that are evenly spread along the chromosomes at an average of 40K were chosen in order to target the desired marker density and genomic distribution. High coverage for parent’s WGS, and the targeted genotyping, yielded a good SNPs panel with almost no missing data or out-filtered SNPs for low quality.

The expected segregation of the selected markers (i.e., the parent’s alleles) is 1:1, meaning the allelic frequency should be 0.5 on average. Indeed, our results, which established the genotyping quality of the SNPs in the F1 population, match this prospect (Fig. 3). The allelic frequency by physical position presentation shows a small number of “ hot spots” that display genomic regions with an unexpected segregation ratio (see black arrow on chromosome 3; Fig. 3), where the frequency of the parent’s allele is significantly deviating from 1:1 ratio. This may indicate that the SNPs in that region could be placed in or near a lethal allele, which its presence in the progeny is not favorable. Such an allele, for example, could be an incompatibility *S* allele^33^.

### Genetic maps construction

Two maps were constructed, each for a different parent (Fig. 5). The map for the U.E.F was denser, with 971 markers divided for all the linkage groups (Table 2, Fig. 5). Both maps were fitted to the eight chromosomes published in the two almond reference genomes (https://www.ncbi.nlm.nih.gov/bioproject/572860; https://www.ncbi.nlm.nih.gov/bioproject/553424). The current U.E.F map density (one marker per ∼0.5 cM) is almost equal to the average recombination frequency (0.475 cM per Mb) (Table 2). Considering this, together with the small population size (n=92), we assume that the recombination rate (i.e., population size) and not the number of markers is the bottleneck for obtaining better mapping resolution.

High co-linearity between the physical and the genetic maps (Fig. 5) demonstrates the reliability of the constructed map. Furthermore, it gives an overview of the genomic distance between the wild almond species, *P. arabica*, and *P. dulcis* cultivar cv. Lauranne, which, remarkably, seems to be quite similar. Moreover, 98% of the sequenced reads of *P. arabica* were mapped to cv. Lauranne reference genome. Although two reference genomes are available for almond, a genetic map is essential for two main reasons. Firstly, genetic map, which relies on recombination frequency of the current population, should represent the most accurate result for marker arrangement of the population compared to the reference genome^34^. In this respect, both genetic maps conform well to the physical data of the cv. Lauranne reference genome. Nonetheless, the few markers which are not correlating could emphasize some chromosomal aberrations as translocation. Secondly, using the genetic map, 38 markers that were delineated as chromosome 0, were linked to LGs by recombination (see yellow dots in Fig. 5).

The F1 population and the character of markers selected, resulted in construction of two maps. SNP markers are mainly di alleles, by choosing SNPs that are heterozygous for one parent and homozygous for the other, we can assure that the markers will segregate, and also to identify the parental origin of each allele. In order to intersect the maps, shared markers are needed. Joining the F1 maps could be achieved by SSR markers that have more than di alleles, as was done in apricot F1 population^35^, or by SNPs that are heterozygous for both parents, as was done in pear^20^.

### SPC genetic mapping

Linkage analysis (i.e., QTL mapping) and GWAS were two approaches used in this study for linking the SPC phenotype and genetic markers (Fig. 6 and Table S3). While GWAS only associates between genomic markers without any data on their specific genomic location, the linkage analysis, which is based on a genetic map connects phenotype to a specific genomic locus / loci^34^.

QTL mapping discovered two significant loci. One major locus with an exceptional LOD score of ∼20 on LG/chromosome 7 was found in the U.E.F map, and another minor (LOD score of ∼4) was detected at the end of LG/chromosome 1 in *P. arabica* map. GWAS revealed both loci. Moreover, the loci identified by the GWAS overlapped those found by QTL analysis. Markers on chromosome 0 also demonstrate significant LOD score (∼20) (Fig. 6, Table S3). Using the genetic map and the QTL analysis enabled the positioning two of these markers to locus 7 QTL (Table S3). Here we demonstrated the importance of integrating the association data, which is based on the physical reference genome and the QTL mapping approach based on the F1 recombination frequency. The overlapping results between the two methods emphasize the unbiased genetic infrastructure and corroborate the mapping data. The data presented identified new genetic loci on the almond genome. To our knowledge, such a mapping effort on the SPC trait has never been done before for any plant, above all, in trees. This strong mapping data is important in order to fully comprehend the physiological role of SPC and identify new genetic components that control photosynthesis in plants. The availability of segregating population and genetic markers highly associated with SPC provides a powerful means to explore this trait.

### Candidate genes

The high-resolution QTL mapping and the robust annotations data sets available, enabled the establishment of a putative list of candidate genes. The list is based primarily on the data from the genetic maps and association studies. The list was filtered for non-synonymous polymorphism in the coding region of the genes between the F1 parents, (Table S4). Based on annotation, the list contains, among others, several genes that are involved in sugar transport (gene ID: Prudu_004403, Prudu_004404, Prudu_004408; Fig. S3). Preliminary results demonstrated a high negative correlation in the F1 population between cork layer (i.e., periderm) development (qualitative 1-5 scale measurements) and the SPC (data not shown). This data suggest that periderm development genes may be involved. For example, *HXXXD*-*type acyl-transferase* and *MYB-family transcription factor* (gene ID: Prudu_018862, Prudu_018912; Fig. S3) genes, which were suggested to be involved in cork synthesis ^36,37^. Interestingly, the *EPIDERMAL PATTERNING FACTOR-like protein 2* (EPF2) gene is a member of this list (Table S3; Prudu_018883). The EPF2 gene is a direct regulator of epidermis cell development and stomatal density^38^. Although the EPF2 gene looks as a promising candidate to control SPC through its role in controlling stomatal density, advanced genetic research should done for establishing the link between the SPC phenotype to this gene. However, this study indicates that the variation of SPC trait in the F1 population is controlled by a small number of genes localized to only two loci in the almond genome. This work sets the stage for further studies aimed to delineate the genetic nature of the SPC trait, define its importance to the tree, and understand how it could be utilized for tree improvement targeted to produce fruit trees adapted to extreme climate. A recent study demonstrated that the cultivated almond breeding lines are highly conserved and are founded only on the cvs. Tuono, Cristomorto, and Nonpareil^39^. This study emphasizes the importance of introduction and utilization of genetic material originating from wild sources. Our current study sets the way for utilizing the wild almond P. arabica as a new source for widening and enriching the current narrow base of almond breeding material.

## Conclusion

This paper is the first to establish a genetic study on mapping the unique stem photosynthetic capability originating from the wild *P. arabica* almond. Here we localized the genetic components that regulate the trait and narrowed the whole ∼240 Mb almond genome towards only two loci, with one major locus spanning only ∼400kb and explaining 67% of the SPC phenotype, which eventually provided a list of 73 candidate genes. Forward genetic approach based on the establishment of a cross-bred population with genetic mapping and GWAS provides a remarkable infrastructure for future introduction of beneficial traits from wild almond origin. This approach is highly efficient for both, study the genetics of important agricultural traits and introducing of new breeding material into highly conserved almond cultivars.

## Material and methods

### Plant material

All trees are growing in the almond orchard in Newe Ya’ar Research Center in the Yizre’el Valley (latitude 34°42’N, longitude 35°11’E, Mediterranean temperate to subtropical climate). The parents of the F1 population, *P. arabica and* the Israeli leading commercial cv. Um el Fahem (U.E.F) are grown at two copies for each, grafted on GF.677 rootstock, and planted in winter 2018. The F1 population (*P. arabica* X U.E.F) contains 92 seedlings that were germinated in the nursery in winter 2017 and replanted in the orchard in winter 2018.

### Gas exchange measurements

Gas exchange measurements were done in the field on one-year old stems (i.e. current year growth) of three years old *P. arabica* and *P. dulcis* (U.E.F) trees, from October 2019 to October 2020. Each month, two reciprocal days were chosen; in each day, four stems per genotype were analyzed (n=8 per month). All measurements were conducted between the hours 8:30-10:30 a.m. (the latest were in winter). When there were leaves on the stems, they were removed two days prior to measurements to eliminate wounding stress effect. Measurements were carried out with the LI-6800 Portable Photosynthesis System (LI-COR Biosciences, USA), using the 6×6-needle chamber, which is compatible with tree branches of 2.5-4.5 mm diameter. The following conditions were held constant in the chamber: photon flux density of 1,200 μmol m^−2^ sec^−1^ (90% red, 10% blue) and CO_2_ reference of 400 PPM was set. Chamber relative humidity, and the temperature held for each month according to the multi-annual average. Gas exchange results were normalized to stem surface area and displayed as net assimilation rates (µmol CO_2_ m^-2^ sec^-1^), transpiration rates (mmol H_2_O m^−2^ sec^−1^), and instantaneous water use efficiency (iWUE; the ratio between net assimilation and transpiration rates).

Gas exchange measurements on the F1 population were conducted in February 2020 for two weeks, while the trees were dormant, between the hours 9:30-11:30 a.m. Four stems were measured (n=4) for each genotype. The measurement protocol was the same as mentioned above. To determine stem respiration rates, stem gas exchange measurements were conducted under the same conditions as described above. Next, the Licor 6800 light source was turned off for ∼2 minutes (for stabilization of ΔCO_2_), and data were recorded. In dark, the net assimilation value represents respiration. Stem respiration rate was recorded under 17°C, 28°C and 34°C.

### Whole genome sequencing (WGS) and SNP calling

DNA extracted from young leaves using the plant/fungi DNA isolation kit (NORGEN BIOTEK CORP, Canada). DNA of *P. arabica* and U.E.F, was sent to Macrogen (Macrogen, Korea) for WGS - Illumina Nova Seq 6000, with targeted coverage of X50 on average, read length of 150 bp with paired-end sequencing. OmicsBox software (version 1.3.11; https://www.biobam.com/omicsbox/) was used for preprocessing the raw-reads based on Trimmomatic^40^ for removing adapters and contamination sequences, trimming low-quality bases, and filtering short and low-quality reads. The cleaned reads were mapped onto the reference genomes: *P*.*dulcis* cv. Lauranne (https://www.ncbi.nlm.nih.gov/bioproject/553424) and *P. dulcis* cv. Texas (https://www.ncbi.nlm.nih.gov/bioproject/572860), using the Burrows-Wheeler Aligner (BWA) software 0.7.12-r1039, with its default parameters^41^. The resulting mapping files were processed using SAMtools/Picard tool (http://broadinstitute.github.io/picard/, version 1.78)^42^; for adding read group information, sorting, marking duplicates, and indexing. Then, the local realignment process for locally realigning reads was performed so that the number of mismatching bases was minimized across all reads using the RealignerTargetCreator and IndelRealigner of the Genome Analysis Toolkit version 3.4-0 (GATK; version http://www.broadinstitute.org/gatk/)^43^. Finally, the variant calling procedure was performed using HaplotypeCaller of the GATK toolkit (https://gatk.broadinstitute.org/hc/en-us) developed by Broad Institute of MIT and Harvard (Cambridge, MA, USA). Only sites with DP (read depth) higher than 20 were further analyzed. SnpEff program^44^ was used to categorize the effects of the variants in the genomes (Table 1, Table S4). The program annotates the variants based on their genomic location (intron, exon, untranslated region, upstream, downstream, splice site, or intergenic regions) including in the Almond GFF file extracted from the NCBI database (GCA_008632915.2). Then it predicts the coding effect such as synonymous or non-synonymous substitution, start or stop codon gains or losses, or frame shifts.

### Population genotyping

Based on WGS of *P. arabica* and U.E.F, a SNP calling was performed in order to select SNPs that will detect polymorphism within the F1 population. The following criteria were set: (1) remove sites with DP lower than 20; (2) an isolated SNP over 100 bp interval; (3) the SNP is unique with no matching on other genomic regions on the reference genome; (4) informative SNPs for the F1 population, that are homozygous for one parent and heterozygous for the other. In addition, SNPs were chosen at intervals of 40 kb along the almond genome (*P. dulcis* cv. Lauranne; https://www.ncbi.nlm.nih.gov/bioproject/553424) to obtain an unbiased representation through the whole chromosomes. Overall, a set of 5,000 markers was selected for genotyping (Fig. 3). The F1 population screening was accomplished by “ targeted SNP Seq” by LGC (LGC Genomics, Germany) for SNPs genotyping.

### Genetic map construction

For generating the genetic map the JoinMap®4.1 software^25^ was used. Cross-pollination population type was used with the code lmxll for markers that were homozygous for the male parents (*P. arabica*) and heterozygous in the female parent (U.E.F), and the code nnxnp for the opposite case. Because there were no common markers (hkxhk) we did not combine the two marker types, and undertook the pseudo test cross method^45^, meaning we separated the markers into two different maps, one map for the U.E.F (where *P. arabica* is homozygous-lmxll code), and one for the *P. arabica* parent (where U.E.F is homozygous nnxnp code). Markers were filtered for three parameters: (1) More than ∼11% missing data; (2) Non-Mendelian segregation (X^2^>6.5, DF=1) (3) Remove markers in similarity of 1.0. The “ Independence LOD” algorithm was used for linkage groups clustering (LOD>8), and the Kosambi’s function was chosen for calculating genetic distance.

### QTL mapping

In order to conduct the QTL analysis we used the Map QTL®5 software^26^. QTLs and their significance were calculated using interval mapping (IM). A QTL was determined as significant when its LOD score was higher than the calculated threshold (1000 permutation at α=0.05), and the QTL spanning was determined by ±1 LOD from the max LOD marker.

### Genome wide association study (GWAS)

Association was calculated by TASSEL 5.2.59^27^. The set of SNPs was filtered; marker discard when missing data was >8.6%, and the allele frequency was set for 0.2<x<0.8 for preventing overestimated impact of rare alleles. The General linear model (GLM) was applied for the phenotypic and genotypic intersect data set to test the association. Threshold for significance result was assessed by 1000 permutation test α=0.05.

### Statistics

All significance tests were done by the statistical software JMP (JMP® PRO 15.0.0 © 2019 SAS Institute Inc.), α=0.05. To test significance when the variance was unequal, a simple T-test was used, and if it was equal, the pooled t-Anova test was performed. Tukey - Kremer’s test was used to analyze variance in the population when the distribution was normal, and the variance inside the groups was equal; when it was not equal or normal, Wilcoxon non-parametric test was used. Broad sense heritability of the SPC was calculated on the F1 (full sibs) by the ‘Rsquare adj’ value, given by a simple Anova test.

## Supporting information

supplementary all tables

## Data availability

All data supporting the results are specified in the manuscript or in the supplementary data. *P. arabica* leaf net CO_2_ assimilation, SPC distribution analysis, and correlation analysis between the SPC and cork development are available from the corresponding author upon reasonable request.

## Acknowledgment

Thanks to Kamel Hatib for planting, grafting and maintaining the experimental orchard. Thanks to Dr. Elad Oren for the close guidance in designing the SNPs panel and processing the genotyping results. The authors thank the generous support of the Leona M. and Harry B. Helmsley Charitable Trust for their help in funding of the current research.

## Author Contributions

HB designed and conducted all experiments and wrote the manuscript. ADF processed and assembled the row sequences of the parent’s DNA, the SNPs calling, analyzing the SNPs effect and all genes’ annotation. IBY and RHB generated the F1 population. TAS, ZA, and HB developed the infrastructure for measuring stem gas exchange. DH is the corresponding author. Designed the experiments, supervised the study, and wrote the manuscript. All authors discussed and commended on the manuscript.

## References

1. Sánchez-Pérez, R., Del Cueto, J., Dicenta, F. & Martínez-Gómez, P. Recent advancements to study flowering time in almond and other Prunus species. Front. Plant Sci. 5, 1–7 (2014).

2. Atkinson, C. J., Brennan, R. M. & Jones, H. G. Declining chilling and its impact on temperate perennial crops. Environ. Exp. Bot. 91, 48–62 (2013).

3. Sperling, O., Silva, L. C. R., Tixier, A., Théroux-Rancourt, G. & Zwieniecki, M. A. Temperature gradients assist carbohydrate allocation within trees. Sci. Rep. 7, 1–10 (2017).

4. Tixier, A. et al. Spring bud growth depends on sugar delivery by xylem and water recirculation by phloem Münch flow in Juglans regia. Planta 246, 495–508 (2017).

5. Fernandez, E. et al. Fruit load in almond spurs define starch and total soluble carbohydrate concentration and therefore their survival and bloom probabilities in the next season. Sci. Hortic. (Amsterdam). 237, 269–276 (2018).

6. Guo, C. et al. Developmental transcriptome profiling uncovered carbon signaling genes associated with almond fruit drop. Sci. Rep. 11, 1–12 (2021).

7. Zwieniecki, M. A., Tixier, A. & Sperling, O. Temperature-assisted redistribution of carbohydrates in trees. Am. J. Bot. 102, 1216–1218 (2015).

8. Roskov, Y. et al. Species 2000 & ITIS Catalogue of Life, 2019 Annual Checklist. Digital resource. http://www.catalogueoflife.org/annual-checklist/2019/details/species/id/939461e1a929360fdd6dc0c65c266d10 (2019) doi:ISSN 2405-884X.

9. Rajabpoor, S., Kiani, S., Sorkheh, K. & Tavakoli, F. Changes induced by osmotic stress in the morphology, biochemistry, physiology, anatomy and stomatal parameters of almond species (Prunus L. spp.) grown in vitro. J. For. Res. 25, 523–534 (2014).

10. Sorkheh, K., Shiran, B., Khodambshi, M., Rouhi, V. & Ercisli, S. In vitro assay of native Iranian almond species (Prunus L. spp.) for drought tolerance. Plant Cell. Tissue Organ Cult. 105, 395–404 (2011).

11. Sorkheh, K. et al. Phenotypic diversity within native Iranian almond (Prunus spp.) species and their breeding potential. Genet. Resour. Crop Evol. 56, 947–961 (2009).

12. Aschan, G. & Pfanz, H. Non-foliar photosynthesis - A strategy of additional carbon acquisition. Flora 198, 81–97 (2003).

13. Nilsen, E. Stem photosynthesis: extent, patterns and rol in plant carbon economy. Plant stems Physiol. Funct. Morphol. 223–240 (1995).

14. De Baerdemaeker, N. J. F., Salomón, R. L., De Roo, L. & Steppe, K. Sugars from woody tissue photosynthesis reduce xylem vulnerability to cavitation. New Phytol. 216, 720–727 (2017).

15. Bloemen, J., Vergeynst, L. L., Overlaet-Michiels, L. & Steppe, K. How important is woody tissue photosynthesis in poplar during drought stress? Trees - Struct. Funct. 30, 63–72 (2016).

16. Sperling, O. et al. Predicting bloom dates by temperature mediated kinetics of carbohydrate metabolism in deciduous trees. Agric. For. Meteorol. 276–277, 107643 (2019).

17. Delplancke, M. et al. Combining conservative and variable markers to infer the evolutionary history of Prunus subgen. Amygdalus s.l. under domestication. Genet. Resour. Crop Evol. 63, 221–234 (2016).

18. Yazbek, M. & Oh, S. H. Peaches and almonds: Phylogeny of Prunus subg. Amygdalus (Rosaceae) based on DNA sequences and morphology. Plant Syst. Evol. 299, 1403–1418 (2013).

19. Miotto, Y. E. et al. Spring is coming: Genetic analyses of the bud break date locus reveal candidate genes from the cold perception pathway to dormancy release in apple (Malus × Domestica borkh.). Front. Plant Sci. 10, (2019).

20. Gabay, G. et al. High-resolution genetic linkage map of European pear (Pyrus communis) and QTL fine-mapping of vegetative budbreak time. BMC Plant Biol. 18, 1–13 (2018).

21. Olukolu, B. A. et al. Genetic linkage mapping for molecular dissection of chilling requirement and budbreak in apricot (Prunus armeniaca L.). Genome 52, 819–828 (2009).

22. Wang, Y., Georgi, L. L., Reighard, G. L., Scorza, R. & Abbott, A. G. Genetic mapping of the evergrowing gene in peach [Prunus persica (L.) Batsch]. J. Hered. 93, 352–358 (2002).

23. Ballester, J., Socias I Company, R., Arus, P. & De Vicente, M. C. Genetic mapping of a major gene delaying blooming time in almond. Plant Breed. 120, 268–270 (2001).

24. Jung, S. et al. 15 years of GDR: New data and functionality in the Genome Database for Rosaceae. Nucleic Acids Res. 47, D1137–D1145 (2019).

25. Van Ooijen, J. W. & J. J. JoinMap®4.1, software for the calculation of genetic linkage maps in experimental populations of diploid species. Kyazma B.V. Wageningen, Netherland. (2013).

26. Van Ooijen, J. W. Map QTL 5, Software for the mapping of quantitative trait loci in experimantal population. Kyazma B.V. Wageningen, Netherlands. (2006).

27. Bradbury, P. J. et al. TASSEL: Software for association mapping of complex traits in diverse samples. Bioinformatics 23, 2633–2635 (2007).

28. Ávila-Lovera, E., Haro, R., Ezcurra, E. & Santiago, L. S. Costs and benefits of photosynthetic stems in desert species from southern California. Funct. Plant Biol. 46, 175–186 (2019).

29. Ávila-Lovera, E., Zerpa, A. J. & Santiago, L. S. Stem photosynthesis and hydraulics are coordinated in desert plant species. New Phytol. 216, 1119–1129 (2017).

30. Khadivi-Khub, A. & Anjam, K. Morphological characterization of Prunus scoparia using multivariate analysis. Plant Syst. Evol. 300, 1361–1372 (2014).

31. Granot, D., David-Schwartz, R. & Kelly, G. Hexose kinases and their role in sugar-sensing and plant development. Front. Plant Sci. 4, 1–17 (2013).

32. Zhang, J. et al. OPEN A new SNP genotyping technology Target SNP-seq and its application in genetic analysis of cucumber varieties. 1–11 (2020) doi:10.1038/s41598-020-62518-6.

33. Gómez, E. M., Dicenta, F., Batlle, I., Romero, A. & Ortega, E. Cross-incompatibility in the cultivated almond (Prunus dulcis): Updating, revision and correction. Sci. Hortic. (Amsterdam). 245, 218–223 (2019).

34. Oren, E. et al. High-density NGS-based map construction and genetic dissection of fruit shape and rind netting in Cucumis melo. Theor. Appl. Genet. 133, 1927–1945 (2020).

35. Hurtado, M. A. et al. Genetic linkage maps of two apricot cultivars (Prunus armeniaca L.), and mapping of PPV (sharka) resistance. Theor. Appl. Genet. 105, 182–191 (2002).

36. Vulavala, V. K. R., Fogelman, E., Faigenboim, A., Shoseyov, O. & Ginzberg, I. The transcriptome of potato tuber phellogen reveals cellular functions of cork cambium and genes involved in periderm formation and maturation. Sci. Rep. 9, 1–14 (2019).

37. Soler, M. et al. A genomic approach to suberin biosynthesis and cork differentiation. Plant Physiol. 144, 419–431 (2007).

38. Hara, K. et al. Epidermal cell density is autoregulated via a secretory peptide, EPIDERMAL PATTERNING FACTOR 2 in arabidopsis leaves. Plant Cell Physiol. 50, 1019–1031 (2009).

39. Howad, W. et al. Pedigree Analysis of 222 Almond Genotypes Reveals Two World Mainstream Breeding Lines Based on Only Three Different Cultivars. (2020).

40. Bolger, A. M., Lohse, M. & Usadel, B. Trimmomatic: A flexible trimmer for Illumina sequence data. Bioinformatics 30, 2114–2120 (2014).

41. Li, H. & Durbin, R. Fast and accurate short read alignment with Burrows-Wheeler transform. Bioinformatics 25, 1754–1760 (2009).

42. Li, H. et al. The Sequence Alignment/Map format and SAMtools. Bioinformatics 25, 2078–2079 (2009).

43. Depristo, M. A. et al. A framework for variation discovery and genotyping using next-generation DNA sequencing data. Nat. Genet. 43, 491–501 (2011).

44. Cingolani, P. et al. A program for annotating and predicting the effects of single nucleotide polymorphisms, SnpEff: SNPs in the genome of Drosophila melanogaster strain w1118; iso-2; iso-3. Fly (Austin). 6, 80–92 (2012).

45. Swinburne, J. & Lindgren, G. Genetic Linkage Maps. Equine Genomics 1137, 11–47 (2013).

