## supplementary all tables for "Revealing the genetic components responsible for the unique photosynthetic stem capability of the wild almond *Prunus arabica* (Olivier) Meikle"

*Supplementary* *information*

**Table S1. Variant’s effect.** Summarized variant’s (SNPs and InDels) effect comparing between *P. arabica* and U.E.F. All data shown analyzed by the SnpEff program (SnpEff 5.0d version). Only SNPs with DP>20 were analyzed.

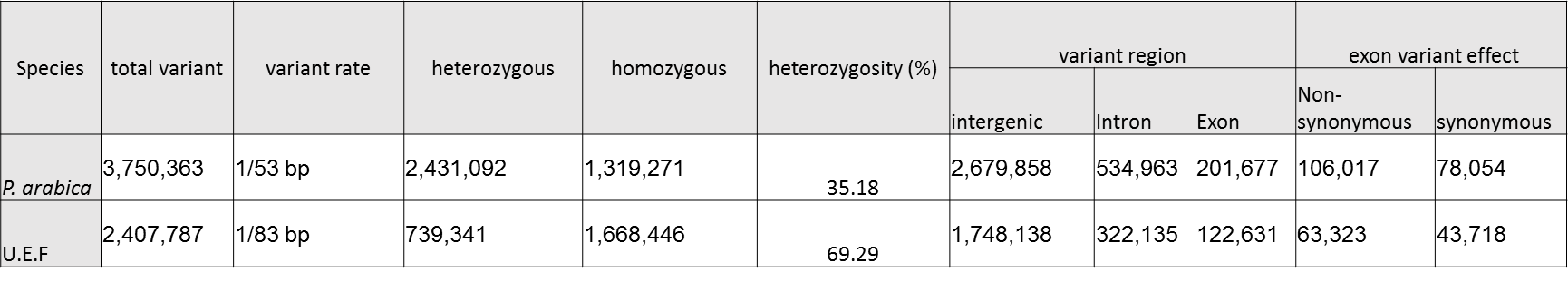

**Table S2**. Genotyping quality parameters for the selected SNPs. Data provided by LGC Genomics (LGC Genomics, Germany).

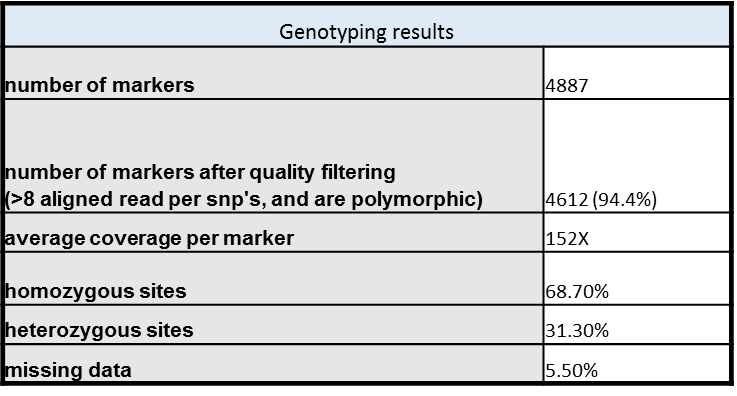

**Table S3. List of highly linked markers, detected by GWAS and QTL mapping, regulating the SPC trait.** Markers highly associated with the QTLs are presented according to their method of detection. Grey markers represent markers from chr-0 which were found by the genetic map on chr-7. The QTL boundaries defined by ±1 LOD or by the significance threshold. Marker ID also represents its physical position.

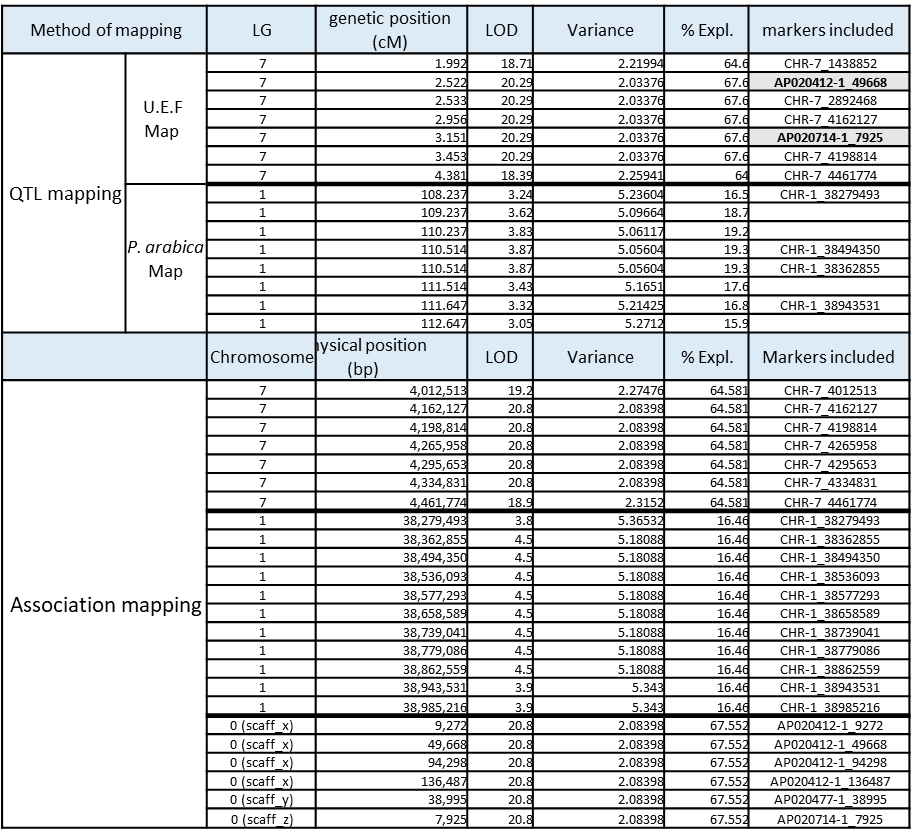

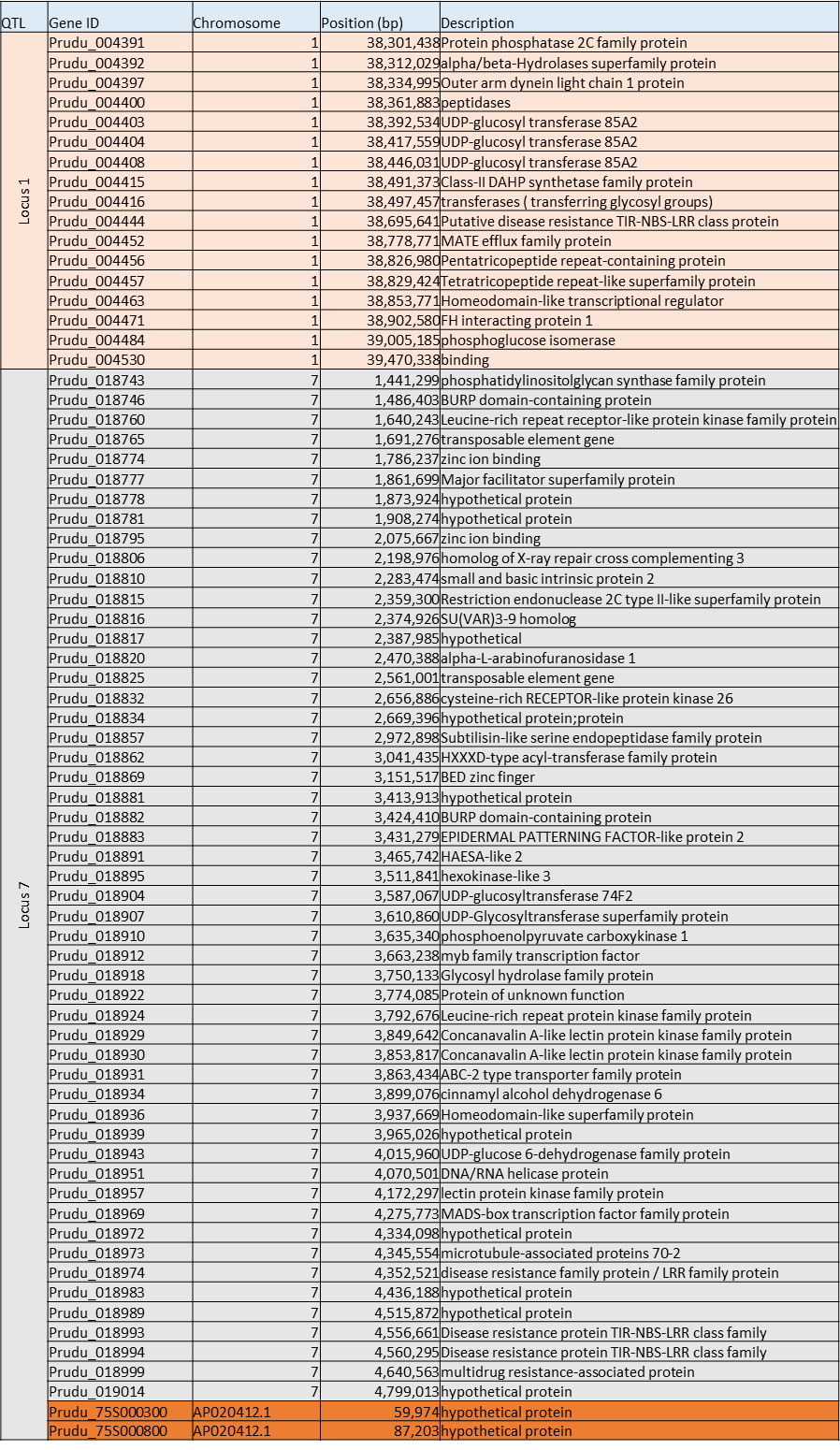

**Table S4. Candidate genes annotation list.** Annotation of the genes located in the DNA region underlined by the two QTL’s according to their physical position. Presented here the candidates list after filtering for genes with only non-synonymous variant located in the coding region. The orange rows represent genes from chr-0 that were genetically mapped to Locus 7.
